# Exploring Anatomical Links Between the Crow’s Nidopallium Caudolaterale and its Song System

**DOI:** 10.1101/2024.07.12.603248

**Authors:** Felix W. Moll, Ylva Kersten, Saskia Erdle, Andreas Nieder

## Abstract

Crows are corvid songbirds that exhibit remarkable cognitive control over their actions, including their vocalizations. They can learn to vocalize on command and the activity of single neurons from the crow’s associative telencephalic structure nidopallium caudolaterale (NCL) is correlated with the execution of this vocal and many non-vocal skilled behaviors. However, it remains unknown if specific anatomical adaptations that directly link the crow NCL to any of the nuclei of the crow’s “song system” exist. To address this issue, we used fluorescent tracers along with histological staining methods (Nissl-, myelin-, and anti tyrosine hydroxylase) to characterize the connectivity of the crow’s NCL in relation to its song system nuclei. We found that the NCL sends dense projections into the dorsal intermediate arcopallium (AID) directly adjacent to and engulfing the robust nucleus of the arcopallium (RA), which is the telencephalic motor output of the song system. Similarly, we demonstrate dense NCL projections into the striatum surrounding the basal ganglia song nucleus “area X”. Both of these descending projections mirror the projections of the nidopallial song nucleus HVC (proper name) into RA and area X, with extremely sparse NCL fibers extending into area X. Furthermore, we characterized the distribution of cells projecting from the lateral part of the magnocellular nucleus of the anterior nidopallium (MAN) to NCL. Notably, a separate medial population of MAN cells projects to HVC. These two sets of connections—MAN to NCL and MAN to HVC—run in parallel but do not overlap. Taken together, our findings support the hypothesis that the NCL is part of a “general motor system” that parallels the song system but exhibits only minimal monosynaptic interconnections with it.

## Introduction

Corvid songbirds (family Corvidae) are vocal learners with large vocal repertoires (Brown and Farabaugh, 1997; Griesser, 2009; Bluff et al., 2010). Their vocalizations play a crucial role in various social interactions, signaling identity and facilitating foraging, pair bonding, or predator mobbing (Griesser, 2009; Kondo et al., 2010; Boeckle and Bugnyar, 2012; Heinrich, 2014). For instance, Siberian jays use their large repertoire of mobbing calls to inform their kin about both predator type and risk posed by the predator (Griesser, 2009). While it is difficult to disentangle the contributions of affect and cognition driving such behaviors in the wild, laboratory studies have recently demonstrated that carrion crows (*Corvus corone*) can exert volitional control over their vocalizations (Brecht et al., 2019; Liao et al., 2024). These crows can vocalize on command, cued by visual or auditory stimuli, and even control the exact number of their vocalizations (Liao et al., 2024). However, the neural pathways controlling this behavior are yet to be determined.

As in other songbirds, the crow brain features an interconnected set of anatomically distinct structures dedicated to skilled vocal production, known as the song system (**Fig. 1**) (Nottebohm et al., 1982; Kersten et al., 2021). This system consists of two pathways (Mooney, 2009), both extending from the vocal premotor nucleus HVC (proper name): (1) the song motor pathway (SMP), driving vocal production in a moment-to-moment fashion via the robust nucleus of the arcopallium (RA) (Hahnloser et al., 2002; Elmaleh et al., 2021; Moll et al., 2023) and (2) the anterior forebrain pathway (AFP), enabling vocal motor exploration and, therefore, vocal learning (Brainard, 2004; Gadagkar et al., 2016; Kojima et al., 2018) HVC receives sensory information from multiple modalities, including auditory, visual, and somatosensory inputs (Wild, 1994; Coleman et al., 2007; Burke et al., 2024). Therefore, it is highly likely that the carrion crow’s song system is involved in producing vocalizations in response to visual or auditory cues (Brecht et al., 2019; Liao et al., 2024). However, functional neural data supporting this hypothesis is currently lacking, as neuronal recordings in vocalizing crows have thus far only been obtained from their NCL (Brecht et al., 2023), a highly integrative multimodal nidopallial area (Kroner and Gunturkun, 1999; Moll and Nieder, 2015, 2017; Kersten et al., 2024). These recordings have shown that the firing rates of single NCL neurons can predict the onset of crow vocalizations following a visual go-cue but not the onset of un-cued vocalizations, suggesting that this premotor activity may be a driver of volitional calls (Brecht et al., 2023). Beyond vocalizations, NCL neurons are also involved in various higher cognitive functions (Rose and Colombo, 2005; Veit and Nieder, 2013; Veit et al., 2014; Hahn and Rose, 2023), of which the representation of magnitudes is a particularly interesting example (Moll and Nieder, 2014; Ditz and Nieder, 2015, 2020; Kirschhock and Nieder, 2022; Wagener and Nieder, 2023) given the crows’ ability to flexibly control the number of their calls (Liao et al., 2024). Collectively, these findings raise the question whether NCL can influence vocal production via direct or indirect connections to any of the song system’s nuclei.

**Fig. 1.**
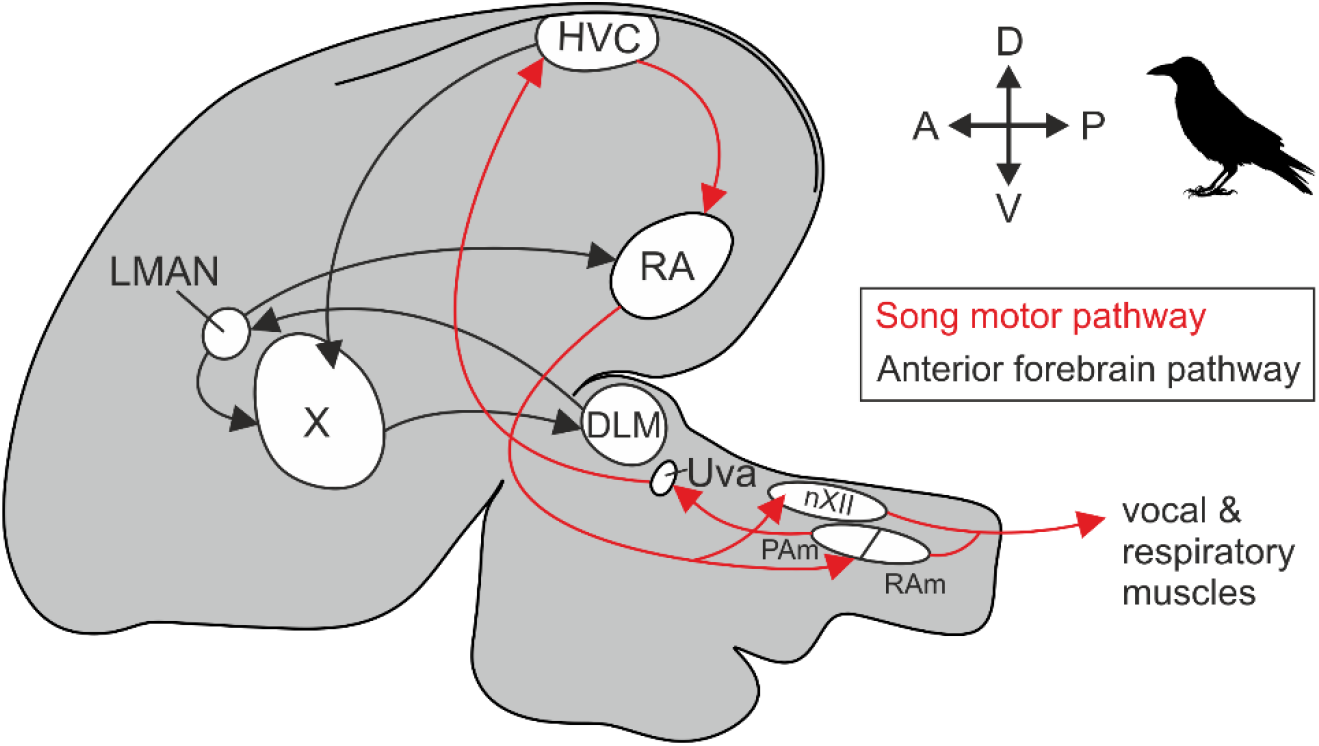
The carrion crow brain exhibits a typical songbird song system. Schematic of the crow brain (side view) showing the position of the main nuclei of the song motor pathway (SMP) and the anterior forebrain pathway (AFP). Based on Kersten et al. (2021). Abbreviations: DLM, nucleus dorsolateralis anterior, pars medialis; HVC, proper name; LMAN, lateral magnocellular nucleus of the anterior nidopallium; nXII, tracheosyringeal nucleus; PAm, nucleus parambiguus; RA, robust nucleus of the arcopallium; RAm, nucleus retroambiguus; Uva, nucleus uvaeformis; X, area X.

Although crow NCL activity correlates with vocal production, direct inter-connections between the NCL and the song system appear to be absent in songbirds such as the zebra finch (Bottjer et al., 2000; Farries, 2001; Paterson and Bottjer, 2017). Instead, NCL seems to be an integral part of a parallel non-vocal ‘general motor system’, which largely mirrors but doesn’t overlap with the brain wide connectivity of the song system (Farries, 2001, 2004; Feenders et al., 2008; Steinemer et al., 2024). As part of this general motor system, NCL likely influences the generation of sensory guided head, beak and neck movements (Knudsen et al., 1995; Feenders et al., 2008; Wild and Krutzfeldt, 2012; Fernandez et al., 2020; Rinnert and Nieder, 2021). However, it is unknown if this general motor system exhibits any specific anatomical adaptations in corvids that could explain the carrion crow’s vocal flexibility (Brecht et al., 2019; Brecht et al., 2023; Liao et al., 2024). We therefore used tract tracing methods to assess direct retrograde and anterograde connections of the NCL in relation to the crow’s well-defined song system nuclei (**Fig. 1**) (Kersten et al., 2021; Kersten et al., 2024).

## Methods

### Animals

We used four hand-raised adult male carrion crows (*Corvus corone*; age range: 8-12 years) obtained from the Institutes’ breeding stock. The crows were kept in spacious aviaries throughout their lives (Hoffmann et al., 2011). All crows had participated in combined behavioral-electrophysiological experiments. The crows’ body weights ranged between 490 and 635 g with rostro-caudal dimensions of the crows’ telencephala between 21.6 and 23.3 mm and brain weights from 7.1 to 7.8 g (measured post-perfusion). All procedures were carried out according to the guidelines for animal experimentation and approved by the responsible national authorities, the Regierungspräsidium Tübingen, Germany.

Brain tissue of all four crows had been previously analyzed for publications focusing on questions different from the scope of the current work, 3 crows for a previous tracing study (cf. **table 1**) (Kersten et al., 2024) and 1 crow for a previously published crow brain atlas (Kersten et al., 2022). For the current study, we prepared additional brain slices from these animals and treated a subset of these slices with immunohistochemical methods. These slices were imaged along with previously mounted slices to thoroughly scan the vicinity of the following song system nuclei: HVC (proper name), the medial and the lateral part of the magnocellular nucleus of the anterior nidopallium (mMAN and lMAN), the tracheosyringeal nucleus (nXII), nucleus parambiguus (PAm), the robust nucleus of the arcopallium (RA), nucleus retroambiguus (RAm), and area X (X). The vicinity of the thalamic nuclei DLM (nucleus dorsolateralis anterior, pars medialis) and Uva (nucleus uvaeformis) has been previously characterized in detail (Kersten et al., 2024).

**Table 1.** Injection protocol for the three crows injected with fluorophore coupled tracers. Each hemisphere was either injected with a dextran amine conjugate (D) or with cholera toxin subunit B (CTB). The fluorophore excitation wavelengths are listed in parentheses.

| Animal | Injections |  | Slicing |  |
| --- | --- | --- | --- | --- |
|  | Left hemisphere | Right hemisphere | Left hemisphere | Right hemisphere |
| Crow #1 | D (555 nm) | D (488 nm) | coronal | coronal |
| Crow #2 | CTB (555 nm) | CTB (488 nm) | sagittal | coronal |
| Crow #3 | CTB (555 nm) | D (488 nm) | sagittal | sagittal |

### Surgical procedures

All surgeries were performed while the animals were under general anesthesia. Crows were anaesthetized with a ketamine/xylazine mixture (50 mg ketamine, 5 mg/kg xylazine initially, supplemented by smaller dosages in regular intervals on demand) and received analgesics (Ditz and Nieder, 2016). During anesthesia, the head was placed in a commercially available stereotactic holder (David Kopf Instruments, Model 1430 Stereotaxic Frame) and ear bars for pigeons (Model # 856 Ear Bars; 20° tapered tip to a 4.8 mm shoulder with a 3 mm dia. by 2 mm long protrusion). A simple beak biting rod was added so that the beak would be held in a 45° angle below the horizontal axis. Based on previously described coordinates (Kersten et al., 2022), NCL was accessed through a small craniotomy centered at 3.5 mm posterior and 11.5 mm lateral relative to the center of the bifurcation of the superior sagittal sinus. This AP position corresponds to ‘AP 4.6’ in our previously published carrion crow brain atlas (Kersten et al., 2022).

### In vivo stereotaxic injections

We used glass pipettes (opening diameter, 20 μm) with an oil-based pressure injection system (Nanoject III, Drummond Scientific) for all tracer injections. Each hemisphere was injected at three medio-lateral injection sites (ML 10.5, 11.8 and 12.5 mm) along one fixed AP value ranging from 3.65 mm to 2.9 mm relative to the center of the bifurcation of the superior sagittal sinus and 1.0 mm below the surface of the brain at a 90° injection angle (i.e., perpendicular to the horizontal plane). We targeted one fixed AP value in a given hemisphere but varied this value at few individual injection sites by a maximum of ±0.2 mm to avoid blood vessel collisions. Individual hemispheres were either injected with cholera toxin subunit B (CTB, Invitrogen, Alexa Fluor 488 or 555 conjugate, C34775 or C34776, respectively) or dextran amine conjugates (10,000 MW, Invitrogen, Alexa Fluor 488 D22910, or fluoro Ruby D1817, 50 mg/ml, diluted in physiological saline solution). We injected 200 nl CTB per site (1.0%, diluted in physiological saline solution; 1 nl pulses at 0.5 Hz with an injection speed of 20 nl/second, total injection time per site: 400 seconds, after the last injection pulse the injection needle was initially left in place for 10 minutes prior to its retraction) and used CTB to analyze patterns of retrograde label as well as anterograde label (Paterson and Bottjer, 2017). Dextran was used to analyze anterograde labeling patterns.

### Histology

Ten days after the tracer injections, crows were injected (i.m.) with 0.5 ml heparin (Braun, 100,000 I.E./10 ml) and a lethal dosage of sodium pentobarbital (Boehringer Ingelheim, Narcoren, 2.5 mL/kg). Subsequently, we perfused the birds with 0.12 M phosphate buffered saline (PBS) including 0.1 % heparin, followed by 4% paraformaldehyde (PFA) in 0.12 M phosphate buffer (PB). The brains were removed from the skull and post fixed in 4% PFA overnight for up to four days. Next, they were sunk in increasing levels of sucrose solution, stopping at a sucrose concentration of 30%. Hemispheres were cut at 50 µm using a cryostat (Leica Biosystems, CM1900) in sagittal or coronal orientation. Slices were collected in PB buffer and stored in antifreeze solution, containing glycerol and ethylene glycol, and stored at -20°C. We mounted series of these slices on SuperFrost™ Ultra Plus object plates (Thermo Fisher) and covered them with Vectashield® antifade mounting medium including DAPI (H-1200 Vector Laboratories). In addition, we Nissl stained a subset of slices as previously described (Kersten et al., 2021). In short, slices were incubated in a warm (55°C) 0.1% cresyl violet solution for 3 minutes, washed in 0.012 M PB, and dehydrated and differentiated in an uprising ethanol series. After immersing them in xylene, they were mounted with Entellan mounting medium (Merck). While most of the crow’s song system nuclei can be identified based on the Nissl stained slices (including HVC, RA, mMAN, and lMANshell), area X and lMANcore cannot be clearly visualized using this staining technique (Kersten et al., 2021). Thus, to identify these nuclei, an antibody staining against tyrosine hydroxylase (TH), the rate limiting enzyme for catecholamine synthesis was performed (Kersten et al., 2021). Slices were treated with 0,3% H_2_O_2_ in PBST (PBS containing 0.2% Triton X-100) to quench endogenous peroxidases, washed and blocked in 5% normal goat serum (Linaris, 69221; Dossenheim Germany, S-1000) and bovine serum albumin (Vector Laboratories, Newark, CA 94560, USA; SP-5050) in PBST. Afterwards, the slices were incubated with the primary anti-TH antibody (mouse anti-tyrosine hydroxylase (TH) antibody, ImmunoStar, Hudson, WI 54016-0488, USA; Cat# 22941, RRID:AB_572268, 1:1000) in PBST and 2,5% NGS for 72 hours with gentle movements at 4°C. After rinsing, sections were treated with the secondary antibody (biotynilated goat-anti-mouse IgG (H+L), Sigma, 82024 Taufkirchen, Germany; SAB 3701068, PRID:AB 2910246, 1:1000) for 2 hours with gentle movements at room temperature. Subsequently, slices were rinsed, incubated in an avidin-biotin complex solution (Elite ABC Kit, PK-6100, Vector Laboratories) and rinsed again. The sections were then developed in 3-3’-diaminobenzidine (DAB) and nickel amplified (DAB Peroxidase Substrate Kit, SK-4100, Vector Laboratories). Afterwards, the stained slices were finally washed and mounted on SuperFrost™ Ultra Plus object plates, dehydrated in ethanol and xylene and mounted in Entellan mounting medium. Furthermore, we used myelin stained slices from a previous data set, to demonstrate the anatomical position of HVC and RA (**Fig. 2a**). For the myelin staining protocol, we incubated mounted crow brain slices in 0,2% gold-chloride solution, fixed the staining result with 2,5% sodium thiosulfate, dehydrated them in ethanol and immersed them in xylene to finally mount and coverslip them in Entellan mounting medium (Kersten et al., 2021, 2022).

**Fig. 2.**
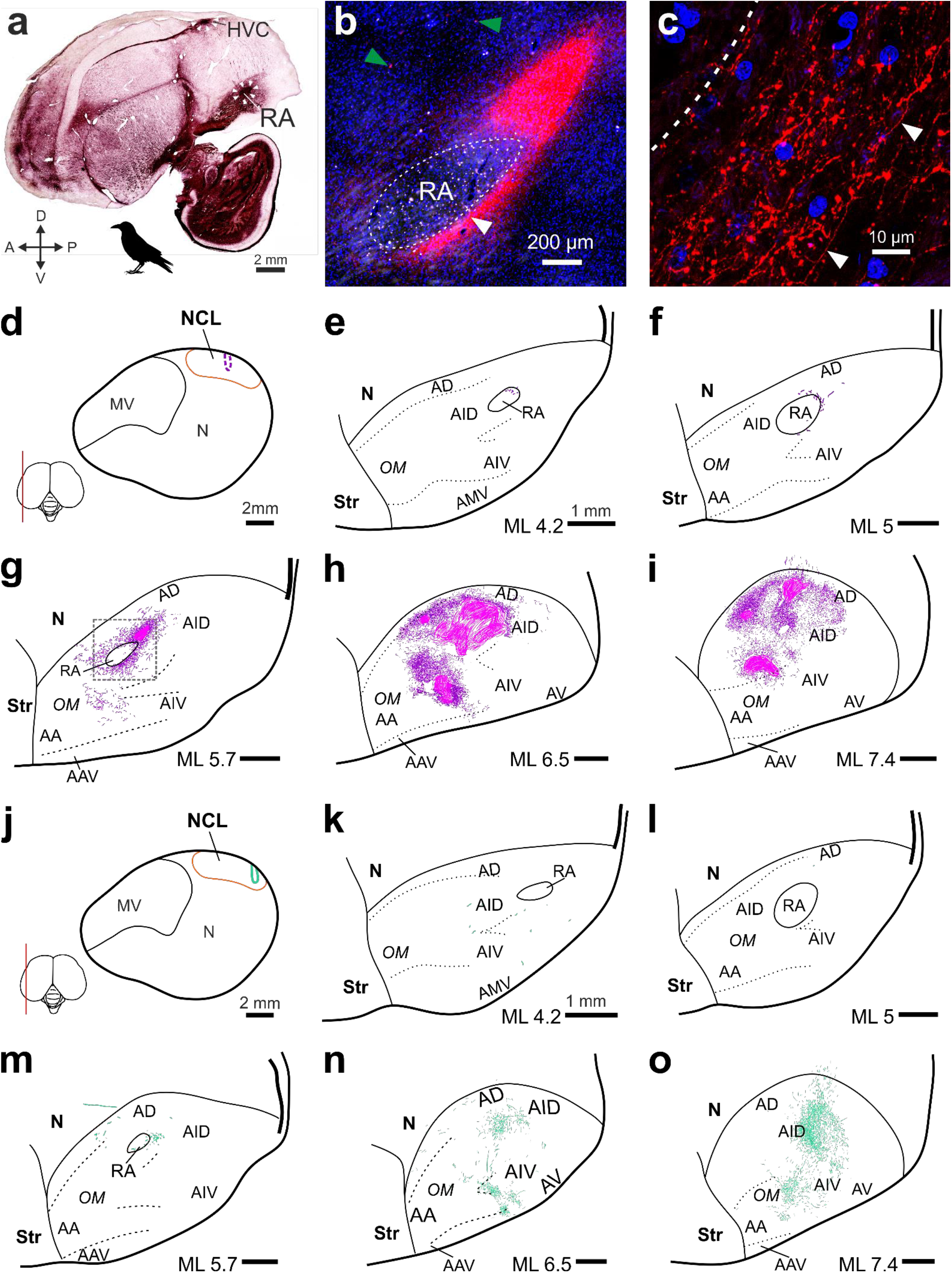
NCL projects to the AID lateral and directly adjacent to RA. **a**, Histological slice of the crow brain at ML 5.0 mm (side view) stained against myelin to show the myelin rich nuclei HVC and RA, which appear as dark ellipsoids indicated by white arrowheads. **b**, Low magnification image of the arcopallium around the lateral RA (dashed outline, ML 5.7 mm). A dense field of CTB labeled NCL fibers (pink) engulfs RA. The position of this image is indicated by the grey dashed rectangle in (**g**). Green arrowheads point at two exemplary retrogradely labeled somata that were absent inside the pink fiber field. Blue: DAPI stained nuclei. White arrowhead indicates the position of the high magnification image shown in (**c**). **c**, Individual anterogradely labeled NCL fibers (red) at the ventral boundary of RA (dashed line). White arrowheads indicate a single continuous fiber running parallel to our imaging plane. Blue: DAPI. **d**, Core of our CTB injection site in NCL at ML 11.45 mm (dashed pink outline) in crow #2, shown in (**b-i**). A conservative estimate of NCL’s extent is indicated by the orange line (cf. Kersten et al. (2022); Kersten et al. (2024)). **e-i**, Arcopallium of the crow also shown in (**b-d**) at increasing ML levels. Magenta, CTB labeled fibers; pink, dense fiber fields (see Methods). **j**, Core of our dextran injection site in NCL at ML 11.9 mm (dashed green outline) in crow #3, shown in (**k-o**). **k-o**, Arcopallium of crow #3. Dextran labeled fibers are shown in green. Abbreviations: A, arcopallium; AA, anterior arcopallium; AAV, anterior ventral arcopallium; AD, dorsal arcopallium; AID, dorsal intermediate arcopallium; AIV, ventral intermediate arcopallium; AMV, medial ventral arcopallium; AV, ventral Arcopallium; OM, occipito-mesencephalic tract; Str, striatum. Scale bars (**e-i**) and (**k-o**), 1 mm. Left is anterior and top dorsal in all panels.

We imaged using a Leica (DMi8) epifluorescence microscope and, to create the high detail fiber images shown in **Fig. 2c** and **3c**, a Leica (Stellaris 8) confocal microscope. Image analysis was performed using Fiji (Schindelin et al., 2012), LAS X (Leica Biosystems) and ZEN (Zen 2.5 lite, blue edition, Carl Zeiss) software. To register the position of individual cells and fiber fragments in a given slice, we first recorded a low magnification overview image, which captured the wider area surrounding RA, X, MAN, or HVC. In the same slice, within the boundaries of the overview image, we recorded tiles of z-stack images at high magnification, which were then matched to the overview image. Using a graphic tablet (Wacom One, Creative Pen Display) and CorelDRAW (Graphics Suite X7), we traced the labeled fibers and registered the labeled cells that could be detected in our high magnification z-stacks. If fiber fields were too dense to discriminate individual fibers, we shaded the area in a different color (**Fig. 2g-i**, pink area). To determine the exact position of individual fibers and cells relative to a given song nucleus and other established brain structures (Kersten et al., 2022), we imaged the same slice with polarized light to identify fiber tracts and laminae. In addition to this information, directly adjacent, Nissl or anti-TH stained slices were used to estimate the exact extent of song system nuclei and their position relative to our registered cells and fibers.

## Results

To test the hypothesis that the crow’s NCL has direct anatomical connections to the song system, we injected fluorescent tracers into the NCL of six hemispheres from three birds (cf. Table 1; tracer spread: ≤ 1 mm around the center of injection sites, cf. Kersten et al. (2024)). Next, we imaged fixed brain slices (50 μm), focusing on all previously identified nuclei of the crow’s song system (see Methods; interslice spacing, ≤ 150 μm) (Kersten et al., 2021). These nuclei were precisely located using previously established crow brain atlases along with histological staining methods (Nissl, myelin, anti TH; see Methods) (Kersten et al., 2021, 2022). We detected labeled fibers and cells in the vicinity of a subset of the crow’s song system nuclei: RA, area X, MAN, and HVC. Furthermore, we detailed the distribution of labeled fibers and cells relative to the boundaries of these nuclei and demonstrate a previously uncharacterized pattern of NCL fibers in the striatum, which distinctly, though not completely, avoid area X.

### NCL densely innervates an arcopallial motor region directly adjacent to RA

NCL projects to the dorsal intermediate arcopallium (AID), located laterally adjacent to RA (**Fig. 2a**) (Bottjer et al., 2000; Mello et al., 2019; Kersten et al., 2024), which drives vocalizations via the brainstem’s syringeal and respiratory nuclei (Wild, 1993; Ashmore et al., 2005; Elmaleh et al., 2021). Therefore, we hypothesized that a subset of NCL-to-AID axons could target AID’s neighbor RA and thereby directly influence flexible vocal production in crows. Injections into NCL resulted in two dense fields of labeled fibers, one restricted to the AID and a second field in the ventral half of the arcopallium, as previously described (Paterson and Bottjer, 2017; Kersten et al., 2024). A subset of fibers in the AID extended medially reaching the direct vicinity of RA in all crows (**Fig. 2b-o**) (n = 6 hemispheres, 3 dextran, 3 CTB, in 3 birds). No labeled somata were detected in AID. Fibers labeled by injections into NCL’s center of mass (n = 2 hemispheres, CTB, in 1 bird) formed terminal fields within AID that extended medially into the neck of the AI (nAID) and RA-cup, two overlapping structures that, together, fully encircle RA (**Fig. 2b-c,e-g**) (Mello et al., 1998; Mandelblat-Cerf et al., 2014; Nevue et al., 2020). Injections posterior to NCL’s center (e.g., **Fig. 2j**) labeled more lateral clusters of fibers in AID, which still sparsely extended medially, reaching RA’s lateral border (**Fig. 2k-o**) (n = 4 hemispheres, 3 dextran, 1 CTB, in 2 birds). However, only few fibers observed near RA extended beyond its estimated border, and even those that did only advanced about 100 µm, despite thorough high-magnification scans of the nucleus (**Fig. 2b-o**). Therefore, our results show that the crow NCL’s direct access to RA is extremely limited, in line with previous studies in other songbird species (Bottjer et al., 2000; Paterson and Bottjer, 2017; Bloomston et al., 2022).

### Area X is scarcely invaded by NCL fibers

NCL’s aforementioned projection to the AID is one of its two large motor related, unidirectional outputs (Kroner and Gunturkun, 1999; Kersten et al., 2024; Steinemer et al., 2024). The second output targets the striatum in crows, as well as in non-oscines such as the pigeon (Kroner and Gunturkun, 1999; Kersten et al., 2024; Steinemer et al., 2024). This projection appears to parallel HVC’s striatal projection to area X (**Fig. 3a**) of the song system’s AFP (**Fig. 1**) (Farries, 2001; Steinemer et al., 2024). However, as noted by Farries (2001, 2004), it remains unclear whether area X is embedded into a terminal field of NCL fibers in the striatum, similar to how RA is embedded within NCL’s terminal field in the dorsal half of the arcopallium (cf **Fig. 2b,g**) (Feenders et al., 2008; Bottjer and Altenau, 2010; Mandelblat-Cerf et al., 2014; Kersten et al., 2024).

**Fig. 3.**
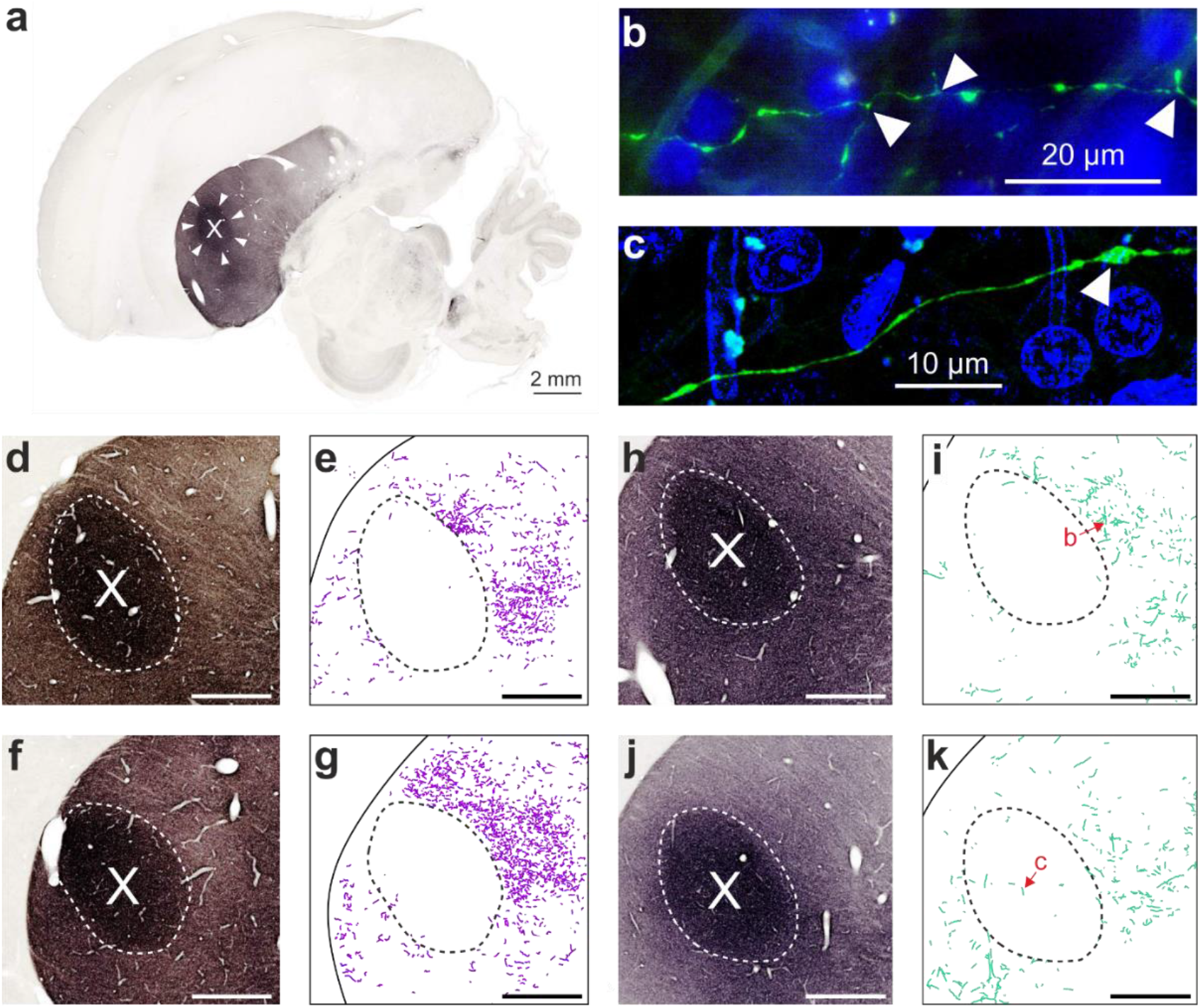
NCL fibers cluster around but rarely invade area X. **a**, Histological slice of the crow brain at ML 4.1 mm (side view) stained against TH, which contrasts area X (white arrowheads) against the striatum around it. **b**, Dextran labeled NCL fiber in the striatum of crow #3 outside of X, position shown in (**i**). Arrowheads indicate short terminal branches. Blue: DAPI. **c**, NCL fiber inside of X, position shown in (**k**). Arrowhead indicates a varicosity, representing a putative en-passant connection. **d-e**, Area X (dashed line) visualized with an anti TH staining in (**d**) and juxtaposed with CTB labeled NCL fibers from the directly adjacent slice in (**e**) (crow #2, ML 3.45 mm). The corresponding injection site is shown in **Fig. 2d. f-g**, Same as (**d-e**) for ML 4.25 mm. **h-i** and **j-k**, Same as (**d-e**) for ML 3.7 mm and 4.3 mm, respectively, but from crow #3 resulting from the dextran injection shown in **Fig. 2j**.

To determine the distribution of labeled NCL fibers within the striatum (e.g., **Fig. 3b-c**) relative to area X, we imaged the vicinity of area X in three birds (n = 6 hemispheres, 3 dextran, 3 CTB). In all birds, we detected many labeled fibers within the medial striatum (ML ∼2 to 6 mm) along with a conspicuous absence of fibers at the estimated position of area X (Kersten et al., 2021). No labeled somata were detected in the striatum. Unlike for RA or HVC, the exact extent of the crow’s area X is difficult to delineate in unstained tissue (Kersten et al., 2021). Therefore, we employed a fine two-step analysis for a representative subset of slices. First, the position of individual fibers was registered at high magnification, blind to the exact boundaries of area X. Additionally, directly adjacent slices were treated with a TH marker (see Methods), which highlights the crow’s area X from the rest of the striatum (e.g., **Fig. 3a**). In a second step, neighboring slices were aligned to determine the border of area X relative to the axonal labeling. This analysis revealed a sharp cut-off of fiber labeling that coincided with the boundaries of area X (**Fig. 3d-k**). Interestingly, fibers outside of area X were arborized, exhibiting short terminal branches (e.g., **Fig. 3b**). In contrast, the few fibers detected in area X were not arborized but still exhibited varicosities, suggesting en-passant contacts (**Fig. 3c**). Therefore, our results show that area X lies *within* the terminal field of NCL’s striatal projection, consistent with the hypothesis of a parallel vocal AFP and a non-vocal “general AFP” (Farries, 2001). Moreover, the presence of few NCL fibers inside area X, exhibiting varicosities, suggests a direct, though sparse, connection.

### LMANshell but not LMANcore and MMAN provide input to NCL

The magnocellular nuclei of the nidopallium (MMAN, LMANcore, and LMANshell) are specific to songbirds and MMAN and LMANcore are considered to be dedicated song system nuclei, projecting to the zebra finch’s HVC and RA, respectively (Vates et al., 1997; Schmidt et al., 2004; Paterson and Bottjer, 2017). Parallel to the MMAN-to-HVC projection, LMANshell (**Fig. 4a**) projects to NCL but is not considered part of the song system, although it might indirectly contribute to vocal learning (Bottjer and Altenau, 2010; Paterson and Bottjer, 2017). Here, we asked whether the crow NCL, unlike the zebra finch NCL, receives direct input from the song system via projections from MMAN or LMANcore.

**Fig. 4.**
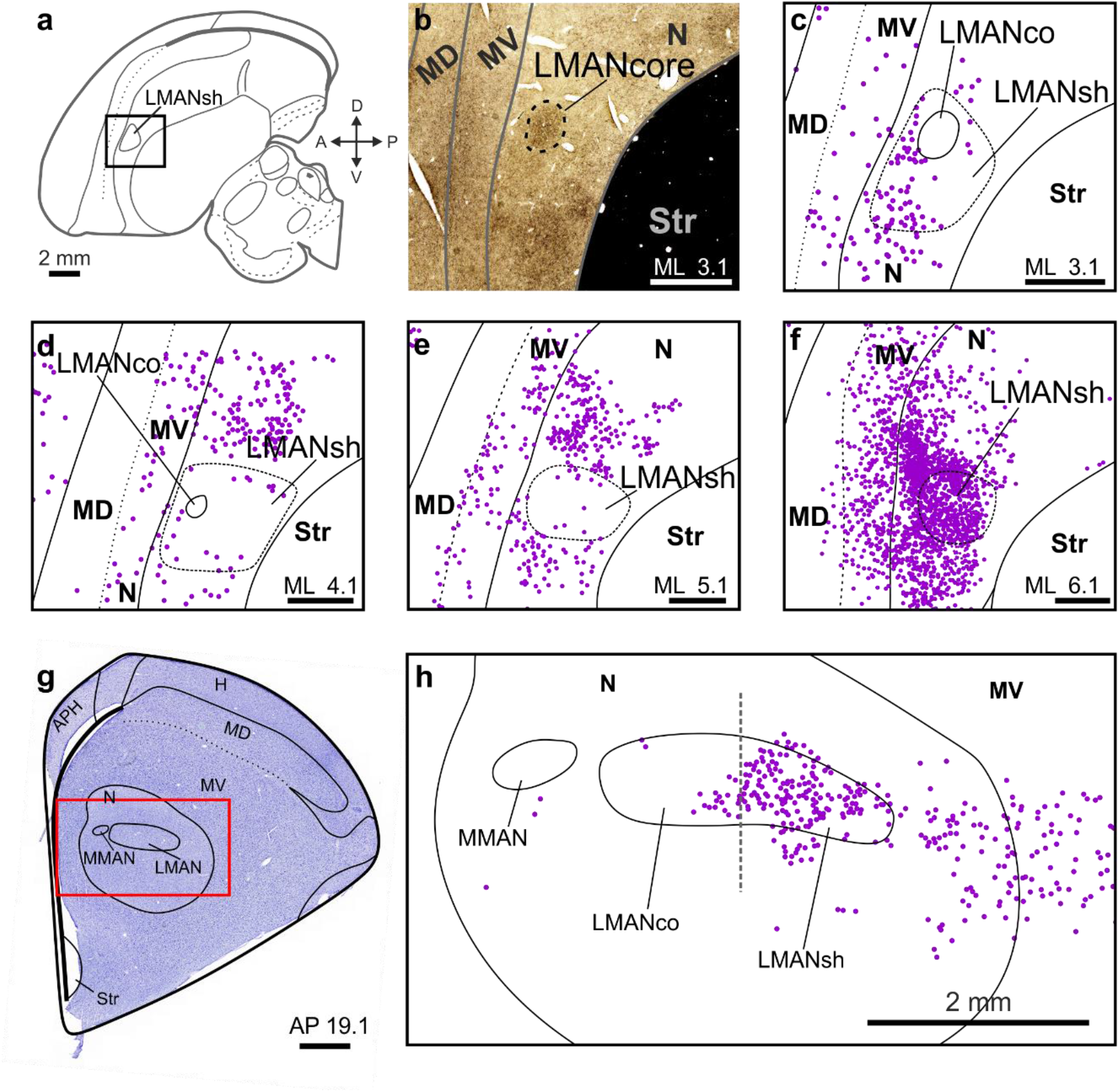
LMANshell projects to NCL. **a**, Crow brain schematic adapted from Kersten et al. (2021) (ML 3.1 mm). The rectangle indicates the position of the histological image shown in (**b**). **b**, TH stained tissue covering the MAN area. **c**, MAN area in the slice directly adjacent to (**b**). Retrogradely labeled somata resulting from a CTB injection into NCL (crow #2, left hemisphere, cf. **Fig. 2** for injection site) are shown as dots (magenta) in relation to anatomical boundaries. The position of LMANcore is estimated based on (**b**). **d-f**, Same as in (**c**) but increasingly lateral to it. **g**, Nissl stained coronal slice (crow #2, right hemisphere). The red rectangle indicates the area from a directly adjacent slice shown in (**h**). **h**, Dots show retrogradely labeled somata (magenta). Although the Nissl stained tissue (**g**) did not reveal the exact extent of LMANcore, we were able to estimate its lateral border (gray dashed line) based TH stained series from the left hemisphere. Abbreviations: APH, hippocampal formation; H, hyperpallium; MD, dorsal mesopallium; MV, ventral mesopallium; N, nidopallium; Str, striatum.

**Fig. 5.**
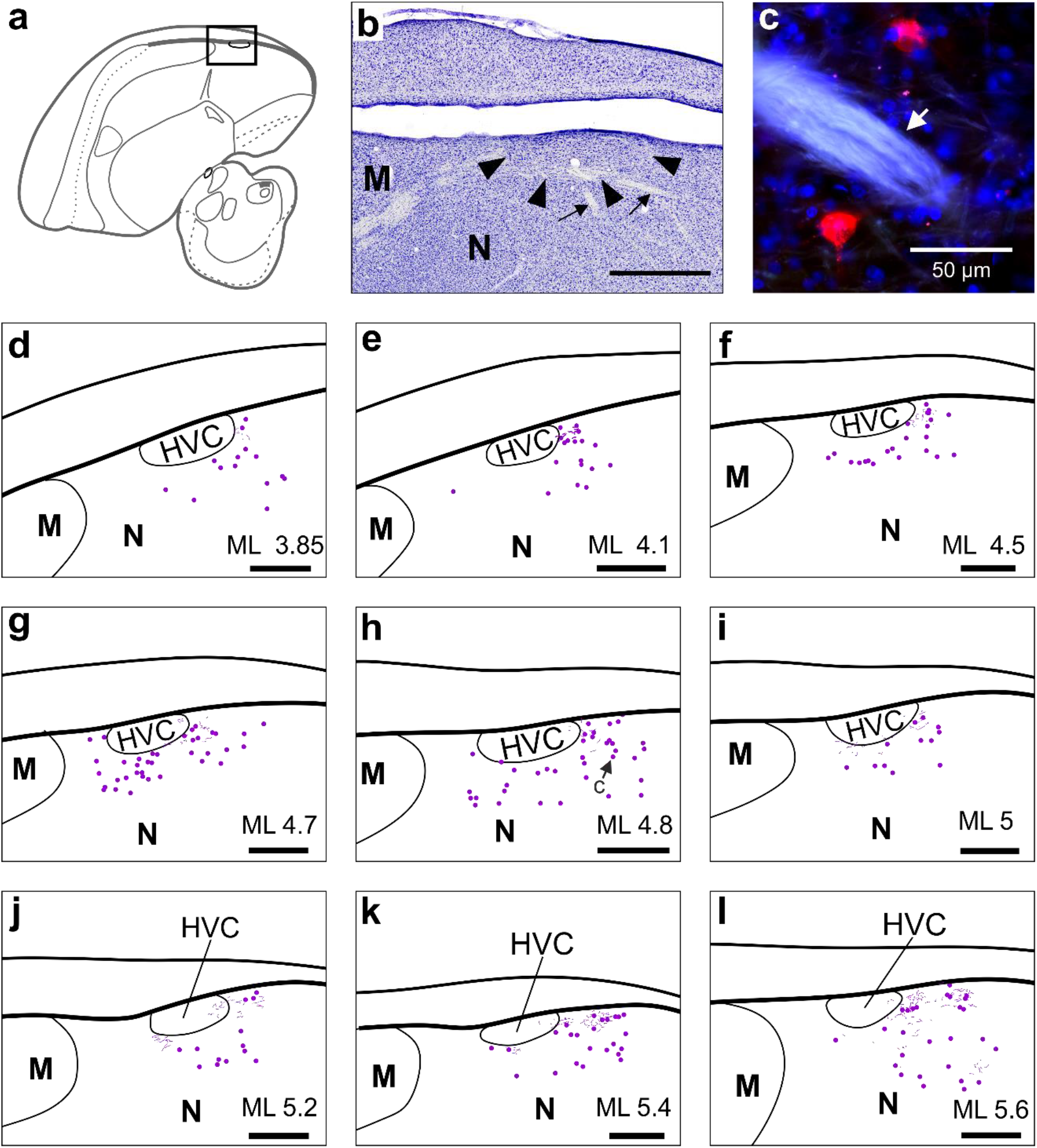
Local NCL connectivity reaches HVC’s vicinity. **a**, Crow brain schematic adapted from (Kersten et al., 2021; Kersten et al., 2024). Rectangle indicates position of histological image shown in (**b**). **b**, Nissl stained tissue showing HVC (arrowheads) and descending HVC fiber bundles (small arrows), ML 4.1 mm. **c**, Two retrogradely labeled somata in the vicinity of HVC, also indicated in (**h**). Injection site shown in **Fig. 2d**. Arrow points at an autofluorescent descending HVC fiber bundle. Blue: DAPI. **d-l**, HVC in the context of labeled cells and fibers (magenta). Abbreviations: M, mesopallium; N, nidopallium.

Somata were densely labeled in the lateral LMANshell and the surrounding intermediate nidopallium (NI), as previously reported (n = 3 hemispheres, CTB, in 2 birds) (Kersten et al., 2024). Although the retrograde labeling efficacy of the fluorophore-coupled dextran was much lower compared to CTB in our crows, we also detected bright dextran labeled cells scattered throughout the lateral LMANshell area (n = 3 hemispheres, dextran, in 2 birds). To determine the distribution of labeled somata relative to LMANcore and MMAN, we registered all somata in the MAN area in a representative subset of slices, blind to the exact positions of LMANcore and MMAN. Additionally, directly adjacent brain slices were stained with anti-TH or Nissl to reveal the extent of LMANcore and MMAN (**Fig. 4b,g**). Neighboring slices were then aligned to determine the boundaries of these nuclei relative to the registered cells (**Fig. 4 b-h**). We did not find labeled cells within the boundaries of MMAN or LMANcore, suggesting that the crow NCL, like the zebra finch NCL, does not receive input from these nuclei (Paterson and Bottjer, 2017).

### HVC-shelf projects to NCL

In crows and other songbirds, NCL is located directly lateral and posterior to the song premotor nucleus HVC in the caudal nidopallium (Paterson and Bottjer, 2017; Kersten et al., 2021, 2022). Therefore, we hypothesized that local NCL projections could invade its neighbor HVC in flexible vocalizers like the crow, even though this connection is absent in songbirds such as the zebra finch (Paterson and Bottjer, 2017).

No labeled fibers or somata were detected in or around HVC in crows that were injected into the posterior half of their NCL (e.g., **Fig. 2j**) (n = 4 hemispheres, 3 dextran, 1 CTB, in 2 birds). However, in one crow injected into NCL’s center of mass (**Fig. 2d**), local labeling extended anterior-medially from the injection site reaching HVC, approximately 7 mm away (n = 2 hemispheres, CTB). In this bird, labeled somata and fibers were loosely clustered around HVC, thus overlapping with the nidopallial area referred to as HVC-shelf (**Fig. 4c-l**) (Mello et al., 1998). Interestingly, few fibers overlapped with the most posterior aspect of HVC (e.g., **Fig. 4j**) but it remains unclear, if these fibers originated from cells at the injection site or, though less likely, if they belonged to close by retrogradely labeled somata. In either case, these sparse fibers could potentially relay NCL signals to HVC cells.

## Discussion

We investigated the anatomical overlap between crow brain structures connected to the NCL and the nuclei of the crow’s song system, revealing a parallel organization of NCL’s and HVC’s input and output connections (**Fig. 6**). Our characterization of NCL fibers in the striatum extends previous findings, demonstrating that the song system’s area X is nested *within* a dense NCL fiber field. In addition, we found few fibers invading area X, which may give NCL direct, though limited, access to this nucleus. Taken together, our findings add to the growing anatomical, cellular, and molecular evidence suggesting that the song system in birds is organized parallel to and likely developed within their general motor system (Farries, 2001; Feenders et al., 2008; Mello et al., 2019; Zemel et al., 2023; Steinemer et al., 2024).

**Figure 6.**
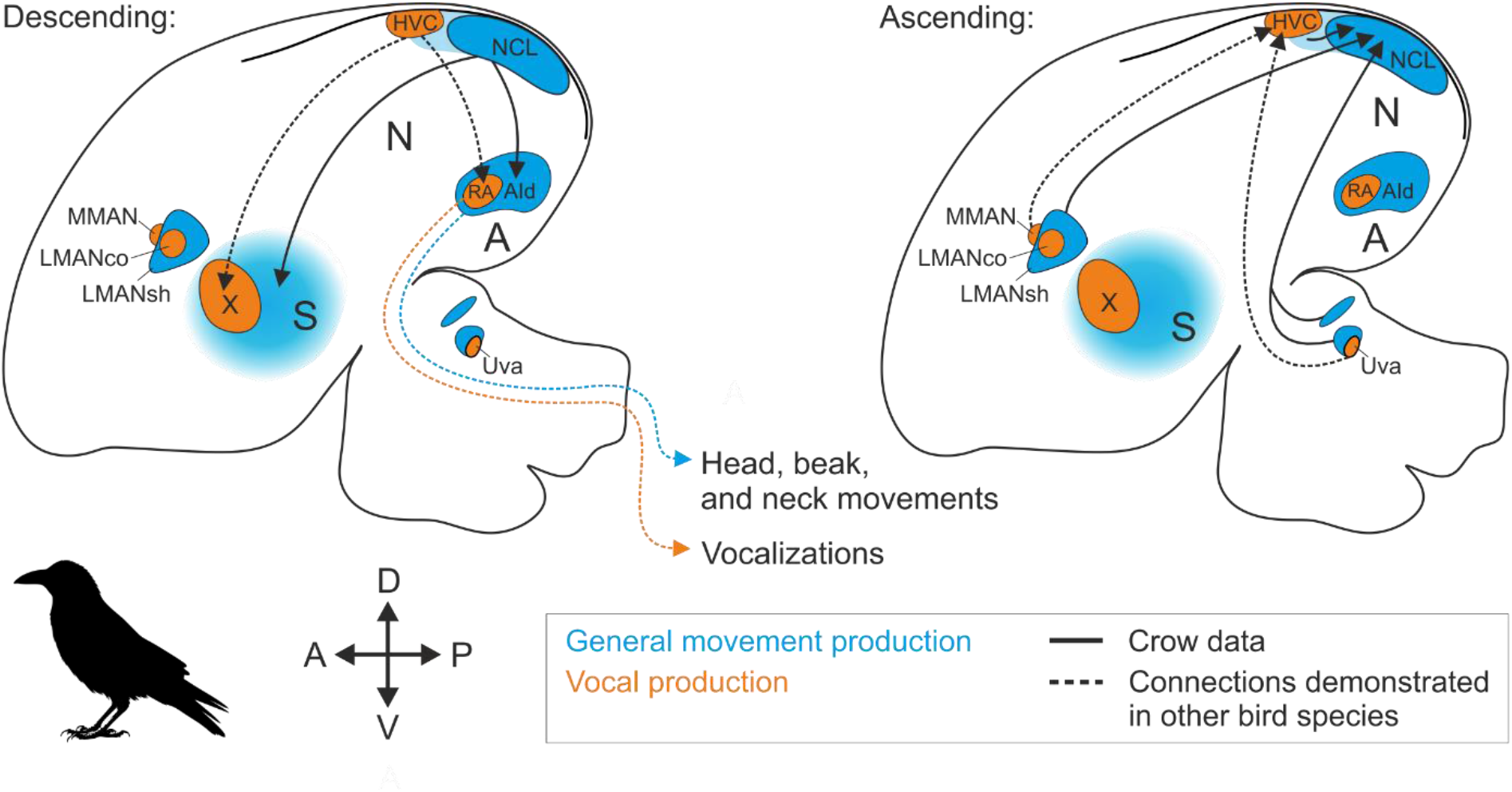
Parallel connections of the vocal premotor nucleus HVC and the general premotor nucleus NCL. Selected structures of the crow’s general motor system (Farries, 2001) are shown in blue, with the light blue area in between HVC and NCL indicating the continuity of NCL’s local connectivity extending to the direct vicinity of HVC (i.e., HVC-shelf). Select nuclei of the crow’s song system are shown in orange (Kersten et al., 2021). Connections between those nuclei (dashed lines) are inferred from and have been demonstrated in other songbird species (Mooney, 2009). Abbreviations: A, arcopallium; AID, dorsal intermediate arcopallium; N, nidopallium; NCL, nidopallium caudolaterale; DLM, nucleus dorsolateralis anterior, pars medialis; HVC, proper name; LMANco and LMANsh, core and shell of the lateral magnocellular nucleus of the anterior nidopallium; MMAN, medial magnocellular nucleus of the anterior nidopallium; RA, robust nucleus of the arcopallium; S, striatum; Uva, nucleus uvaeformis; X, area X.

Skilled vocal production depends on the song system’s telencephalic output nucleus RA (Ashmore et al., 2005; Benichov et al., 2016; Elmaleh et al., 2021). Therefore, we asked whether the NCL has anatomical access to this essential vocal motor hub. Previous studies on the arcopallial projections of the caudal nidopallium (NC) in zebra or Bengalese finches have shown dense terminal fields outside and very few fibers within RA’s boundaries after injections into the dorsal NC directly medial to HVC (Bloomston et al., 2022), into HVC-shelf (Mello et al., 1998; Mandelblat-Cerf et al., 2014), or into NCL (Bottjer et al., 2000; Paterson and Bottjer, 2017). While NC fibers within RA could be a result of tracer spill-over into HVC in some of these cases, we can exclude this possibility for our crows as the tracer spread less than 1 mm around the injection sites, which were located at least 7 mm from HVC. This potential difference might explain the complete lack of fibers that invaded RA’s center in our crows, which are otherwise a consistent result of injections into HVC-shelf (Mello et al., 1998; Mandelblat-Cerf et al., 2014). Thus, we did not find evidence that the crow NCL has greater anatomical access to RA than in other songbirds.

In contrast to RA, we did observe scarce NCL fibers, exhibiting putative en-passant connections well inside the body of area X. Still, area X primarily stood out from the surrounding medial striatum due to a lack of fibers, compared to the dense NCL projections that surrounded it. As these projections have not been previously detailed in songbirds (Bottjer et al., 2000; Bottjer and Altenau, 2010), it will be important to determine whether sparse NCL fibers inside of area X are exclusive to corvids or a general feature among songbirds. Another difference to the arcopallium was that the extent of NCL projections inside the striatum was less topographic (Paterson and Bottjer, 2017), a finding that was also seen in pigeons after injections into different aspects of their NCL (Steinemer et al., 2024). Interestingly, injections into the pigeons auditory aspect of the NCL resulted in a rather limited patch in the medial striatum, at a relative position that is very similar to the position of area X in songbirds (Steinemer et al., 2024). Collectively, these observations suggest that area X is integrated into a pre-existing pallial motor pathway (Steinemer et al., 2024), as proposed by Farries (2001) and referred to as the “general AFP”.

Parallels to the connectivity of the song system’s HVC do not only exist at the output side of the crow NCL (**Fig. 6a**). When considering NCL’s inputs (**Fig. 6b**), projections from LMANshell are particularly dense, surrounding the song system’s LMANcore that is devoid of NCL projecting cells in both our current work and previous work in the zebra finch (Paterson and Bottjer, 2017). LMANcore projects to RA in the zebra finch and this projection is paralleled by one subpopulation of LMANshell neurons projecting to AID (Bottjer et al., 2000; Paterson and Bottjer, 2017). A separate LMANshell subpopulation projects to NCL, parallel to the projection of MMAN, which selectively sends input to HVC (**Fig. 6b**) (Johnson et al., 1995; Vates et al., 1997; Bottjer et al., 2000; Koparkar et al., 2024). Thus, two major projections of the MAN complex (to NCL and AID) are mirrored by the two song system specific projections of MMAN and LMANcore (to HVC and RA, respectively).

Additional parallel input reaches HVC and NCL from the thalamus (**Fig. 6b**) (Nottebohm et al., 1982; Wild, 1994; Wild and Gaede, 2016; Kersten et al., 2024). In the case of HVC, the thalamic multisensory nucleus uvaeformis (Uva) relays feedback from the respiratory system and this signal has a role in starting vocal elements such as calls or individual syllables within a song sequence (Moll and Long, 2023; Burke et al., 2024). In the crow, a distinct cluster of cells adjacent to Uva projects to NCL (Kersten et al., 2024) and it is an interesting open question whether this thalamic input is involved in starting elements of non-vocal movement sequences.

In line with previous work in the zebra finch, our data suggests that NCL is not monosynaptically connected to the body of its neighbor HVC (Bottjer et al., 2000; Paterson and Bottjer, 2017). However, NCL’s local connectivity extended medially invading the direct periphery of HVC (i.e., ‘HVC-shelf’), which, in turn, sends very sparse projections into HVC (Mello et al., 1998), consistent with our observation of few fiber fragments overlapping with the most posterior aspect of HVC. Even though they are sparse, these fibers could potentially route NCL signals to HVC and future studies will determine whether this input is relevant for flexible vocal production.

Considering their anatomical position and connectivity, several related hypotheses on the evolutionary origin of the song system have suggested that HVC might be a specialized part of NCL that could have diverged from NCL via pathway duplication (Farries, 2001; Feenders et al., 2008; Chakraborty and Jarvis, 2015). In these frameworks, the ‘general motor system’ is seen as the evolutionary precursor of the entire song system (Farries, 2004; Feenders et al., 2008; Chakraborty and Jarvis, 2015). This scenario has functional implications for structures like the HVC-shelf or its downstream target RA-cup, which have been interpreted as accessory structures to the song system (Mello et al., 1998; Bottjer and Altenau, 2010). However, as aptly stated by Michael A. Farries (2001): *“These accessory structures may be nothing more than the oscine equivalents of the non-oscine regions* [i.e., the general motor system] *from which the song system emerged, structures that were literally pushed aside by the growth of specialized subdomains within them that became the song system.”*. Following this argument, the vocal domain and the ‘general motor’ domain of the songbird brain may function largely independently.

## Conclusion

While sparse local interactions cannot be excluded at the various hubs of the parallel song and general motor system, NCL’s direct anatomical access to the song system seems to be limited. Instead, NCL is likely primarily involved in controlling movements of the head neck and jaw (Knudsen et al., 1995; Farries, 2001; Feenders et al., 2008; Wild and Krutzfeldt, 2012; Mandelblat-Cerf et al., 2014; Fernandez et al., 2020; Rinnert and Nieder, 2021). This, however, doesn’t exclude the possibility that the crows’ ability to vocalize on command critically depends on NCL (Brecht et al., 2019; Brecht et al., 2023; Liao et al., 2024), a hypothesis that can be tested in future studies. Beyond vocal production, the general motor system of crows and other smart birds offers an opportunity to explore the understudied neurobiology of skilled, non-vocal avian behaviors such as nest building (Hall et al., 2015) or skilled food extraction, including tool use (Cristol and Switzer, 1999; Striedter, 2013; Cabrera-Alvarez and Clayton, 2020).

